# Embedding wear assessment in musculoskeletal simulation: A proof-of-concept application to total hip arthroplasty

**DOI:** 10.64898/2026.08.31.748225

**Authors:** Andrea Di Pietro, Lorenza Mattei, Francesca Di Puccio

## Abstract

Predicting wear in artificial joints requires integrating joint dynamics, contact mechanics and progressive surface evolution, yet these processes are often treated separately. In total hip arthroplasty (THA), finite-element approaches remain the reference standard, but they are computationally demanding and usually rely on boundary conditions from independent musculoskeletal (MSK) models, hindering consistent coupling and feedback between wear progression and movement dynamics. As a single-subject proof of concept, we present a computational framework that embeds wear estimation within forward MSK simulations through OpenSim–MATLAB integration. Contact variables are computed using an elastic-foundation formulation, and wear is updated through the Archard law, enabling cyclic prediction of contact mechanics and surface evolution within a single workflow at practical computational cost. The framework was evaluated in one subject with right THA during five activities of daily living and numerically benchmarked against finite-element simulations. A long-term walking analysis of 4 million cycles was also performed to assess geometry updating. Across tasks, peak contact pressures remained within 7% of finite-element predictions. Linear wear depth and volumetric loss showed maximum deviations of 16% and 13%, respectively. Accounting for progressive geometry changes yielded a maximum wear depth about 31% lower than linear extrapolation. These preliminary results support the framework’s computational feasibility and numerical consistency for the tested case; nevertheless, multi-subject evaluation is required before broader predictive or clinical use.

## 1 Introduction

The long-term durability of total hip arthroplasty (THA) remains limited by wear-induced particle generation, which promotes aseptic loosening [1–3], recognized as one of the main causes of implant failure [4]. In this context, as the average age of THA recipients decreases, and their post-operative activity levels increase [5,6], predictive evaluation of wear under patient- and lifestyle-specific conditions has become essential for improving implant longevity [7–9]. Instrumented-implant datasets, such as OrthoLoad [9], have demonstrated substantial inter-patient and inter-activity variability in hip contact forces, further emphasizing the importance of individualized biomechanical assessment [7–9].

The finite element method (FEM) remains the benchmark approach for simulating contact mechanics and wear in THA, providing high spatial resolution of stresses, contact pressures, and surface damage [10–14]. However, its practical implementation is often labour-intensive and technically demanding. Each simulation requires detailed meshing and contact definitions, as well as careful calibration of numerous parameters to solve nonlinear analyses and ensure numerical stability and reliable predictions [15]. Long-term wear predictions typically rely on accelerated strategies implemented by ad hoc user subroutines. Moreover, most commercial FEM platforms operate within license-based environments, which restrict user access to solver source code [16].

In current practice, boundary conditions applied to FEM wear analyses are typically derived from MSK simulations, which provide the resultant joint forces and rigid-body kinematics acting at the hip during representative motor tasks [17–19]. While this MSK-to-FEM coupling enables realistic load application, it also introduces additional computational complexity and potential inconsistency between solvers.

On the other hand, MSK modelling, as implemented in OpenSim, offers a flexible and fully open computational environment for the dynamic analysis of human movement [20]. Its architecture enables straightforward modification of model structures, contact geometries, and model parameters, while supporting subject-specific simulations of experimentally observed motor tasks [21]. The primary approach currently available in OpenSim to enable contact is the Elastic Foundation method (EFM), which handles penetration between general 3D surfaces represented by triangular meshes [22]. Since EFM relies on a compliant, penetration-based formulation, the contact response depends nonlinearly on both the selected contact parameters and the local penetration depth. Moreover, increasing stiffness to limit penetration may compromise numerical stability and increase implementation challenges, while both the predicted penetration and the computational cost remain highly sensitive to stiffness and mesh density [23].

A key limitation of the OpenSim implementation of the EFM is that spatially resolved contact variables (e.g., local pressure and penetration fields), which are required by wear laws (e.g., Archard-type formulations) are not available as standard simulation outputs [24], despite contact being computed within Simbody [25]. To the best of the authors’ knowledge, a comprehensive, integrated workflow capable of integrating joint contact mechanics with wear prediction remains unavailable, representing a significant limitation for long-term implant performance assessment and motivating the development of a dedicated computational solution.

The present work addresses this gap by introducing an MSK-embedded wear (MEW) framework that enables the direct computation of contact and wear variables for soft-on-hard joint prostheses without relying on external FEM solvers, thereby preserving wear progression within a consistent multibody framework. The framework is examined in a single-subject proof-of-concept THA case across multiple activities of daily living to: (i) demonstrate workflow feasibility and identify task-specific loading conditions; (ii) benchmark contact and wear outputs against FEM simulations under matched assumptions; and (iii) illustrate long-term wear evolution during walking. The study is intended to assess computational feasibility and numerical consistency rather than population-level generalizability or clinical predictive validity.

## 2 Methods

### 2.1 Case study: subject and motor tasks

This single-subject computational study was designed as a proof of concept. It used data from a participant with a right total hip replacement who had provided informed consent before performing several activities of daily living. Experimental data were obtained from the dataset published by Lunn et al. [26] within the LLJ European project [27] and refer to the subject LLJ_140, a male with a body mass of 64 kg and a stature of 172 cm. Five motor tasks representative of common lower-limb activities were analyzed, encompassing both locomotor and non-locomotor actions: normal walking, fast walking, lunge and stair negotiation (i.e. stairs ascent and descent). Walking is prescribed in standard wear-testing protocols (e.g., ISO 14242); stair negotiation was included to represent a more demanding locomotor task. In contrast, the lunge task is biomechanically representative of movements such as obstacle negotiation or certain sport-related actions, including those observed in tennis. A detailed description of the experimental protocol and task execution is provided by Lunn et al. [26]. For this proof-of-concept analysis, one representative trial per task was selected, consistent with the selection adopted in [17].

### 2.2 MSK models

Two MSK models were defined in OpenSim to simulate the case study: a baseline model with natural hip joint and a modified model with right THA, as depicted in Figures 1(a) and (b), respectively. They were used in different phases of the workflow, as detailed in Sec. 2.3.

**Figure 1.**
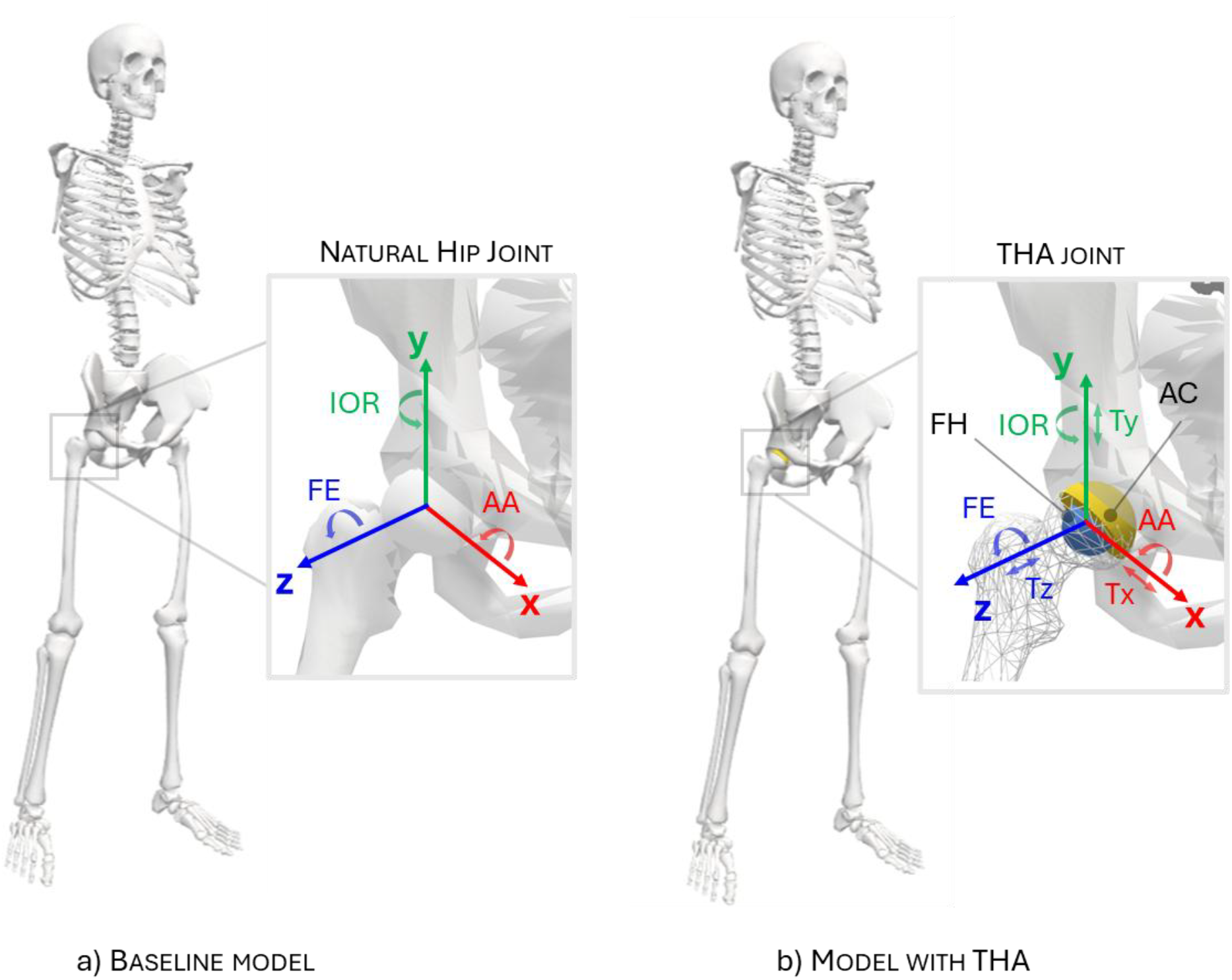
(a) Baseline model (Gait2392) with an ideal spherical hip joint and three rotational DOFs (FE, AA, IOR). (b) Model with THA having six DOFs (adding translations Tx, Ty, Tz) and acetabular cup–femoral head (AC-FH) contact elements.

#### 2.2.1 Baseline model

A musculoskeletal model with 23 degrees of freedom (dofs) and 92 musculotendon actuators (Gait23-92 [28]) was used and scaled to the participant’s anthropometric dimensions (baseline model) using the Automated Scaling Tool [29]. The model represents the trunk and lower limbs, while the inertial effects of the arms are lumped with those of the torso. The 23 dofs are distributed as follows: 6 describe pelvic translation and rotation relative to the ground; 3 define the lumbar joint and the hip ball-and -socket articulations; and one dof is assigned to each of the knee, ankle and subtalar joints. The model is actuated by 92 Hill-type muscle–tendon units, with 43 actuators assigned to each lower limb and 6 to the trunk.

As in most multibody models, the hip joint was modelled as a spherical joint through the combination of three sequential rotational joints following an ISB-compliant rotation sequence: flexion/extension (FE) about the mediolateral axis, abduction/adduction (AA) about the anteroposterior axis and internal/external rotation (IOR) about the longitudinal axis (Figure 1(a)).

#### 2.2.2 Model with THA

The baseline model was subsequently modified to incorporate a right total hip arthroplasty, including an acetabular cup (AC) and femoral head (FH), rigidly fixed to the pelvis and femur, respectively. The hip joint was modelled with 6 dofs, adding to the baseline model rotational dofs, three translational dofs (Figure 1(b)).

##### a) Implant geometry and materials

The THA joint was represented as a ball-and-socket joint with spherical contact surfaces. As no implant-specific information was available in the dataset published by Lunn et al.[26], a metal-on-plastic THA configuration with a nominal head diameter of 28 mm was assumed. Accordingly, the FH was modelled as a sphere of radius 14 mm whilst the AC was defined as a hemispherical cup having internal radius of 14.08 mm (resulting in a clearance of 80 μm) and a thickness *h* of 8 mm.

The cup was considered in anatomical position, with the north pole direction inclined by 45° with respect to the longitudinal axis.

The AC was assumed to be composed of ultra-high-molecular-weight polyethylene (UHMWPE) and was modelled as a linear elastic material with a Young’s modulus (*E*) of 500 MPa and a Poisson’s ratio (*v*) of 0.4. The FH, assumed to be made of structural steel and significantly stiffer than the cup, was therefore modelled as a rigid body.

##### b) Mesh

The contact surfaces were meshed in Meshlab [30]. The resulting surface meshes were exported as STL files and subsequently imported into OpenSim as both bodies and contact geometries (Figure 1). The meshes consisted of uniformly distributed triangular elements [30], with a mean element edge length of 0.675 ± 0.049 mm. The inner surface of AC was discretised with approximately 6300 triangles, interfacing with approximately 12500 triangles representing the FH surface. Mesh resolution was determined on the basis of a mesh sensitivity analysis, in which the influence of AC and FH surface discretization on contact variables, such as maximum contact pressure, was evaluated. Walking conditions were used as the reference loading scenario for this analysis. Starting from a baseline discretization, four progressively finer and four progressively coarser mesh resolutions for both AC and FH were generated (as reported in Table 1), with element dimensions ranging from 0.538 to 0.864 mm.

**Table 1.** Summary of the mesh sensitivity analysis for the MEW.

|  |  |  |  |  |  |  |  |  |
| --- | --- | --- | --- | --- | --- | --- | --- | --- |
| # AC elements | 4452 | 4884 | 5336 | 5808 | 6812 | 7344 | 7896 | 8468 |
| # FH elements | 8833 | 9670 | 10565 | 11500 | 13487 | 14541 | 15002 | 16766 |
| Element size (mm) | 0.803 ± 0.059 | 0.767 ± 0.056 | 0.734 ± 0.053 | 0.703 ± 0.051 | 0.649 ± 0.047 | 0.625 ± 0.045 | 0.603 ± 0.043 | 0.582 ± 0.042 |
| Δ max wear depth (%) | -3.57 | -2.14 | -1.43 | -0.71 | 0.71 | 0.71 | 1.43 | 1.43 |
| Δ V loss (%) | -4.9 | -3.54 | -2.18 | -1.09 | 1.09 | 1.91 | 2.72 | 3.27 |
| Δ Max Pressure (%) | -2.05 | -0.21 | -0.31 | -1.23 | 0.31 | 0.1 | 0.21 | 1.23 |
| Δ CPU time (%) | -34.67 | -18.64 | -1.4 | 4.98 | 28.02 | 41.38 | 28.36 | 32.49 |

##### c) Contact modelling

The interaction between the contact surfaces of the AC and FH was implemented using the EFM available in OpenSim [22]. In this formulation, each triangular facet of the contact mesh is associated with a nonlinear spring–damper element acting at its centroid along the facet normal direction. For each triangle in contact, the normal contact pressure *p*is defined as:

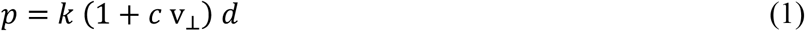

where *d* is the normal penetration at the triangle centroid, v_⊥_ is the normal component of the relative velocity between the contacting bodies at the same location, and *k* and *c* the contact stiffness and damping coefficients, respectively. The stiffness was computed according to the following expression [22]:

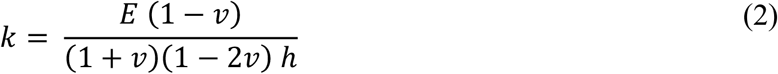

yielding a value of 1.13 10^11^ N/m^3,^ given the above reported material properties and cup thickness *h*. A finely tuned dissipation coefficient *c* of 3 s/m was also included in the contact behaviour to damp elastic oscillations during the dynamic equilibrium.

The contact was assumed as frictionless, as friction effects have been shown not to significantly influence contact variables in metal-on-plastic THA [31].

### 2.3 MSK-embedded wear method

#### 2.3.1 Simulation strategy

Using the baseline model with idealized mechanical joints, a standard inverse MSK workflow estimates joint kinematics from marker trajectories, computes joint moments via inverse dynamics, and resolves muscle forces (e.g., through static optimization). Joint reaction forces are then obtained from dynamic equilibrium using the estimated muscle and external loads.

When force-based contact is included in the THA model, the inverse-dynamics workflow used for the baseline model cannot be applied directly to its unconstrained translational DOFs. Contact forces instead depend on the evolving system states and regulate joint motion without imposing explicit kinematic constraints. Therefore, forward-dynamics-based analyses, namely Computed Muscle Control (CMC) and Forward Dynamics (FD), were used for this formulation.

However, performing CMC directly on the model with THA can be computationally expensive and may exhibit convergence difficulties due to the increased hip joint mobility during the motion tracking and the non-linearity introduced by contact. Conversely, a standard FD simulation is based on open-loop time integration and can suffer from numerical drift and instability, making convergence difficult to achieve. To address these limitations, this study combined the two approaches: the CMC solution obtained from the baseline model (initialization phase) was used to initialize the cyclical FD simulation of the THA model and to provide muscle-control inputs. FD simulation was run over short time windows to avoid drift, and each window was re-initialized from the corresponding CMC-derived states to improve numerical stability and convergence.

A schematic overview of the simulation strategy is shown in Figure 2 and is described in detail in the following sections. Briefly, an initialization phase, performed in the OpenSim environment and adopting the baseline model, provided the state and muscle-control inputs required by the wear computation. The latter was based on a combined OpenSim–MATLAB analysis using the model with THA. Finally, our developed MSK-embedded wear (MEW) method was numerically benchmarked against the FEM approach.

**Figure 2.**
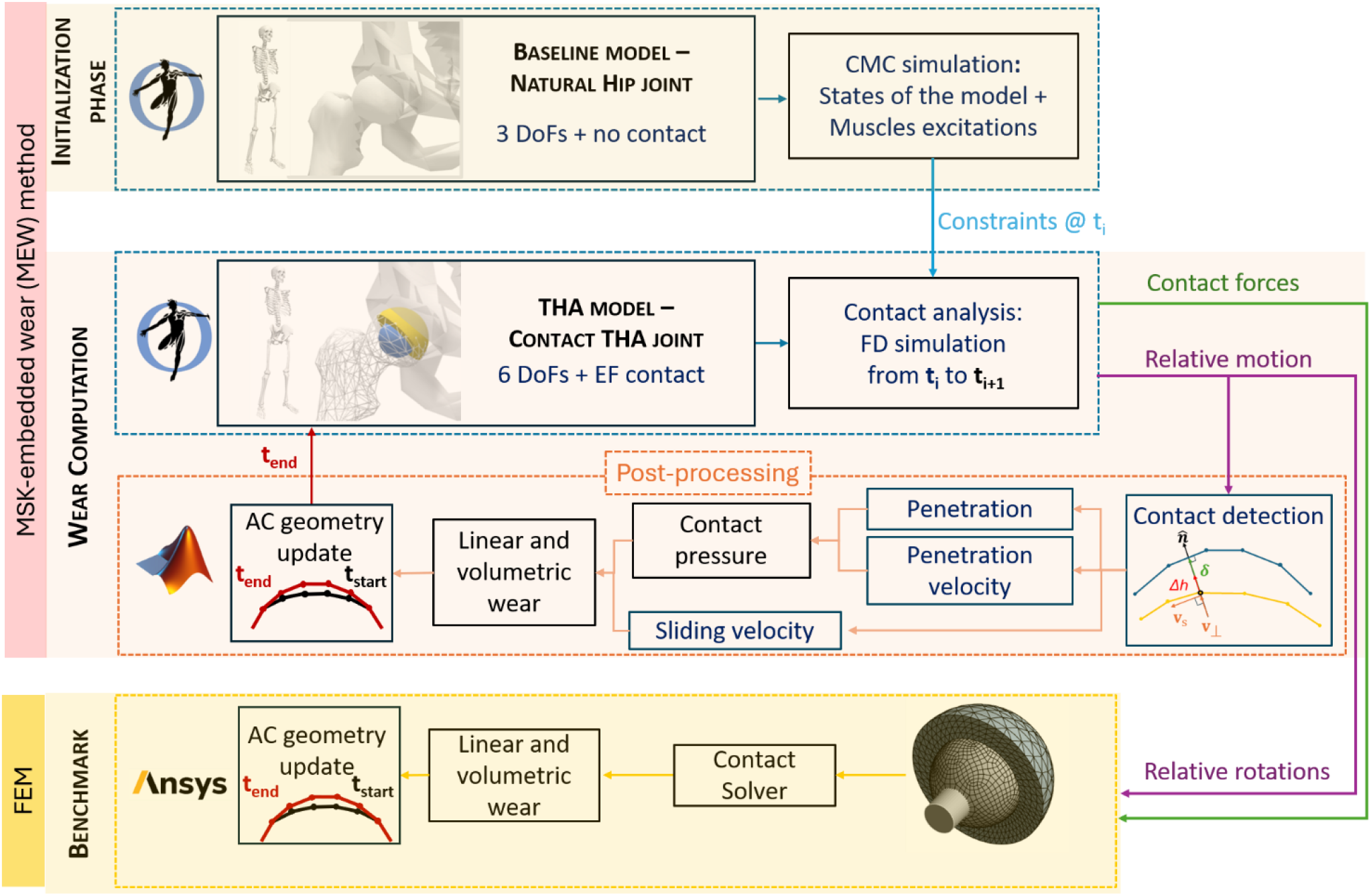
Workflow of the study: the MSK-embedded wear (MEW) method comprises both the initialization phase and the wear computation phase using MATLAB and OpenSim environments. MEW results are numerically benchmarked against finite element method (FEM) simulations in Ansys.

#### 2.3.2 Initialization phase

During the initialization phase, the CMC analysis was performed over the entire motor task to compute (i) model states, and (ii) muscle excitations which provided the state initialization and muscle-control inputs for the FD simulations (see the top of Figure 2). Model states describe the instantaneous dynamic configuration of the system, including joint positions and velocities (generalized coordinates and speeds) as well as muscle fibre lengths and muscle activation states. Muscle excitations, in contrast, represent the neural control inputs to the muscles and govern the time evolution of muscle activation through activation dynamics.

To ensure that the simulated motion matched the experimentally observed joint kinematics, we used the fast target tracking formulation. In this approach, the algorithm finds muscle excitations that reproduce the desired joint accelerations while keeping the overall control effort as small as possible (i.e., minimizing the sum of squared controls). To improve numerical stability and facilitate convergence, small reserve and residual actuators were included; their optimal forces were kept minimal so that they contributed as little as possible to the resulting joint loads.

#### 2.3.3 Wear computation

The computation of wear was performed using the model with THA; as illustrated at the centre of Figure 2, it is based on the cyclic repetition of the following steps: (a) the FD simulation to solve the motion equations and (b) the MATLAB post-processing routine that computes contact and wear estimates and updates the geometry.

##### a) Forward dynamics simulation

At each iteration, an FD simulation is performed in OpenSim over the current time window (Δt=0.01 s), initializing the system with the CMC-derived states and applying the corresponding muscle excitations as controls (Figure 2, central box, dashed blue). During this simulation, the Elastic foundation contact forces contribute to the system’s dynamic equilibrium as a result of the interaction between the contact geometries (i.e., AC and FH) [32]. According to the EFM, contact forces arise from surface penetration driven by the three hip translational dofs and are computed using eq.(1) [25].

##### b)MATLAB post-processing routine

In OpenSim v4.5, which was used in this study, the contact variables required for wear calculations could not be extracted directly [24]. Therefore, contact pressure and sliding velocity were computed using a dedicated post-processing routine implemented in MATLAB® R2025a (The MathWorks, Inc., Natick, MA, USA).

The routine takes as input the STL files of the FH and AC mesh surfaces, together with the OpenSim kinematic outputs describing their relative motion. The latter are expressed as the hip 6 coordinates, i.e., three relative rotation angles and three translations, along with the corresponding angular and linear velocities. The routine provides, as output:

- Contact variables
- Wear assessment (i.e. wear depth and wear volume)
- Worn geometry update

##### Contact variables

Contact pressure is calculated according to the EFM in Equation (1), which requires the penetration depth and its time derivative at contact points. First, the relative pose of the contact pair is reconstructed following the procedure described in [33], mapping the FH mesh into the AC reference frame. Subsequently, contact detection is performed by identifying the AC mesh vertices lying within the FH volume [34]. For each detected contact vertex, the penetration *δ* is computed as the minimum distance between the vertex itself and the FH surface mesh [34], as illustrated in Figure 3. The penetration velocity is obtained by processing the relative velocity between the AC and FH bodies. For each AC contact vertex, the relative velocity vector is decomposed into a normal (v_⊥_) component representing the penetration velocity, and a tangential (v_*s*_) component along the nearest FH facet, representing the sliding velocity.

**Figure 3.**
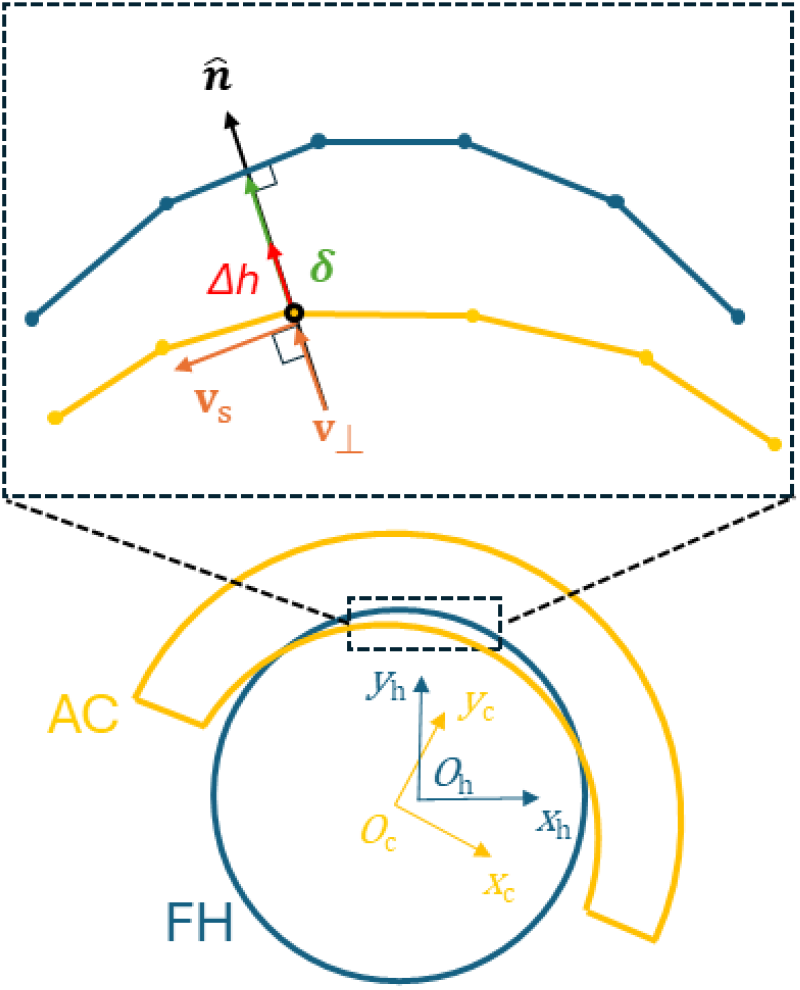
Representation of the post-processing contact and wear variables in the MSK-embedded wear (MEW) method.

##### Wear assessment

The routine implements the Archard wear law [10], because it is widely used in this application [14,35,36]. Additionally, unilateral wear of the AC was assumed, neglecting damage of the much harder metallic component. However, other wear laws, including bilateral formulations, could also be implemented. In its general local and instantaneous form [37] the Archard wear law states that:

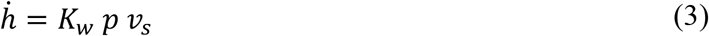

where *K*_*w*_is the wear coefficient (assumed equal to 1.066 ·10^-15^ m^2^/N for metal-on-plastic THA [38]), *p* the contact pressure and *v*_*s*_ the sliding velocity. Numerical implementation of this equation required a temporal discretization, yielding the linear wear increment Δ*h* at each vertex of the AC over the time increment Δ*t* (from *t*_*i*_ to *t*_*i*+1_), considering the average values of both the contact pressure and the sliding velocity:

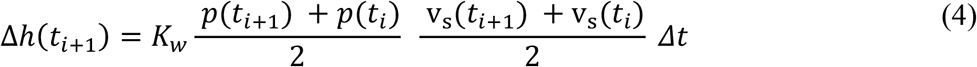

From these increments the evolution of the linear wear map and its maximum value can be easily obtained for the analyzed task. Another important element for wear assessment is the worn volume, which is calculated using the divergence theorem [39].

##### Worn geometry update

Since wear is assumed to occur only on the AC, its STL geometry is updated in the OpenSim model, within the MATLAB routine, only at the end of the motor-task wear computation; specifically, the AC contact vertices are displaced along the surface normal 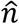 by Δ*h*. Updating the geometry during the task analysis would be unnecessary, as the resulting changes are negligible and do not affect the contact variables.

#### 2.3.4 Accelerated strategy for long-term wear calculation

Once wear variables are calculated for a single task execution, two options are possible to evaluate long-term wear, i.e. wear after N task repetitions: a) simply multiply all wear results by *N*, under a linear-behaviour assumption, which is a conservative assumption; b) repeat the MEW procedure with an updated geometry of the AC to include wear effects. This second option is one of the main novelties of the proposed framework. When the number of repetitions is high, as in the examined application where it is on the order of millions of cycles, it is convenient to adopt an accelerated strategy, as proposed in the literature [40,41], to reduce computational time, combining the above options a) and b). In this case, the total number of repetitions N is divided into k steps of n = N/k cycles each. The wear increment calculated in each step is multiplied by n, and the geometry is then updated accordingly to approximate the cumulative effect of repeated loading. As a proof-of-concept long-term assessment, a wear simulation of 4 million cycles (Mc) was performed for the walking trial, divided into 16 steps, each one corresponding to 250,000 cycles. Notably, the updated STL file is therefore used in the OpenSim model with THA in the subsequent wear step, so that contact pressure and wear are computed on the progressively modified surfaces.

### 2.4 Numerical benchmarking against FEM

For each investigated motor task, a dedicated finite element model was built in Ansys® Workbench (Ansys Inc., Release R2024a) and used as a numerical benchmark for the proposed MEW (Figure 2). Numerical benchmarking was performed by comparing the contact pressure distributions and wear maps predicted (at one wear cycle) by the proposed MEW against the corresponding FEM results.

#### a) Implant geometry and materials

The finite element model shared the same implant geometry (including the 14 mm FH radius) and materials described in Sec. 2.2.2. Accordingly, the femoral head was simulated as a rigid body.

#### b) Mesh

Mesh density was selected via a sensitivity study. Convergence was assumed when further refinement altered peak contact pressure and maximum wear depth by less than 5%. The final discretization consisted of 1236 triangular faces for the AC (modelled with Tet10 elements) and 1332 quadrilateral faces for the FH (modelled with Quad8 elements), with a characteristic edge length of 1.5 mm for both components.

#### c) Boundary conditions

The boundary conditions were prescribed by imposing the task-specific components of contact loads, i.e. joint reactions, and AC-FH relative rotations obtained from the THA model (see Figures 2 and 4). Specifically, both the loading and the kinematic conditions were applied to the FH while the three translational DOFs were left unconstrained and governed by the applied loads. On the other hand, the backing surface of the AC was fully constrained.

**Figure 4.**
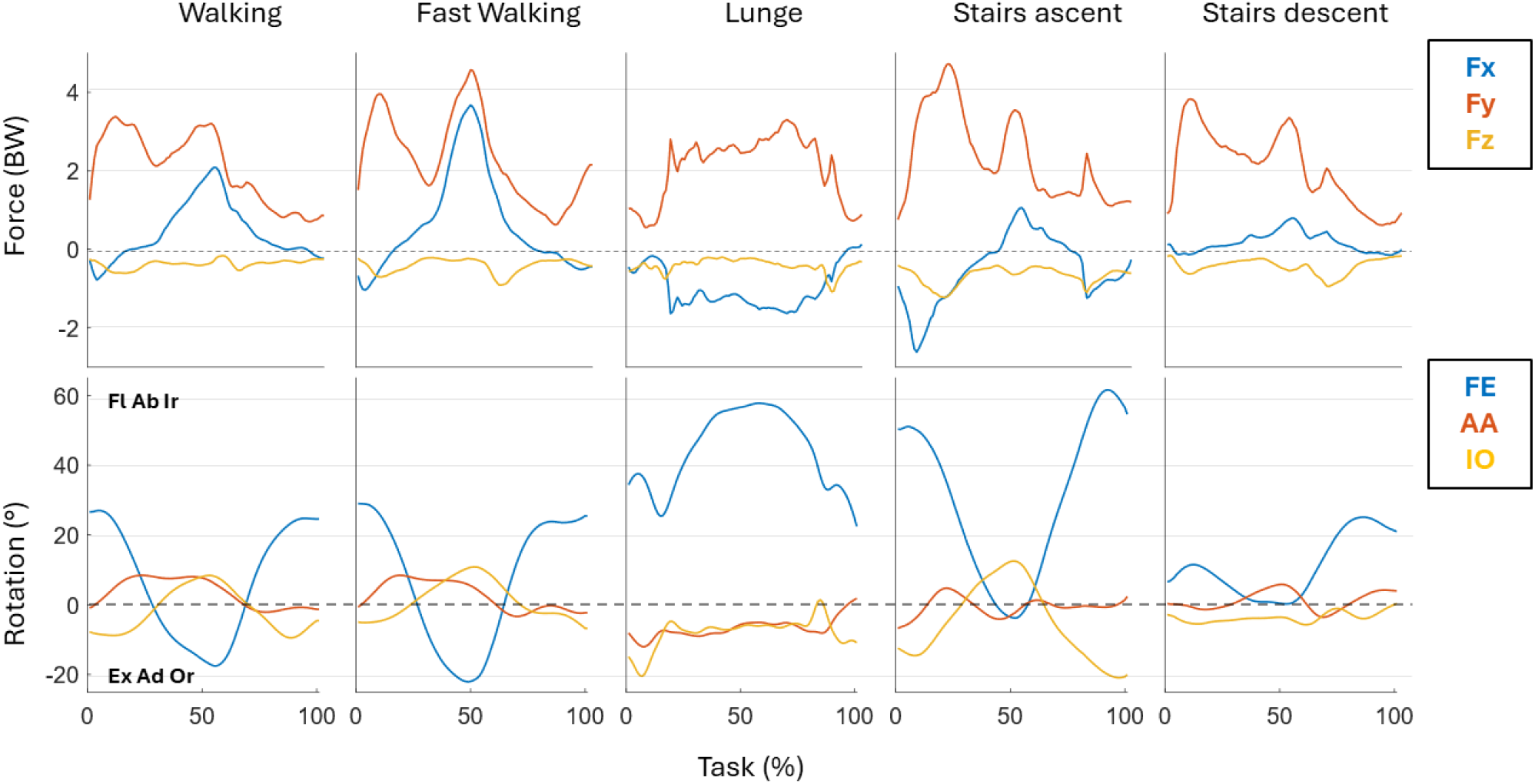
Boundary conditions retrieved from the FD simulations and applied to the FEM simulation.

#### d) Contact modeling

The AC and FH surfaces were defined as contact and target surfaces, respectively, and discretized using CONTA174 and TARGE170 elements. Contact was solved using an Augmented Lagrange formulation with a nodal-normal to target contact detection scheme. Penetration tolerance and normal contact stiffness were left as *program-controlled*, while stiffness updating (*update stiffness*) was enabled at every iteration to improve numerical stability and accuracy. As assumed by the MEW, the contact was simulated as frictionless [31].

#### e) Wear modelling

Wear was modelled in ANSYS Workbench–Mechanical using Contact Surface Wear with the Archard law (TB, WEAR–ARCD via APDL command snippet), assuming dry contact and constant wear coefficient (as reported in Section 2.3.3), with wear applied through geometry updating by moving the contact nodes along the contact normal.

## 3 Results

### 3.1 Mesh sensitivity analysis

The mesh sensitivity analysis for the model with THA, carried out on the single walking trial, indicated a systematic dependence of the output metrics on surface discretization (Table 1). Over the investigated range (4452–8468 elements), maximum wear depth exhibited a modest monotonic increase with mesh density, spanning −3.57% to +1.43% relative to the baseline discretization (6300 elements). A clearer trend was observed for volume loss, which increased monotonically with mesh refinement, with variations of −4.90% to +3.27%. By contrast, maximum pressure showed limited sensitivity, remaining within −2.05% to +1.23%. Computational cost, however, was more strongly affected by mesh density: CPU time varied from −34.67% up to +41.38%, confirming the expected increase in runtime with finer discretization.

### 3.2 Musculoskeletal analysis: hip joint kinematics and reaction forces

Among the outputs of the MEW method were the AC-FH contact forces and the corresponding relative kinematics during task execution, which were implemented as boundary conditions for the FEM simulations (Figure 4). Since the latter were obtained through a tracking algorithm, their accuracy was evaluated by quantifying the kinematic tracking errors during both the CMC and FD procedures. As these errors remained within the recommended optimality range [42], the experimentally observed motion was reproduced within the adopted tracking-error criteria (Supplementary Table S1). The simulated kinematics showed a systematic behaviour across tasks: flexion–extension occurred for most of the relative motion, with the largest excursion during stairs ascent (nearly 60°), whereas stairs descent exhibited the lowest sagittal-plane demand (approximately 25°). The lunge, in contrast, was characterized by a sustained flexed posture throughout the movement.

A consistent feature across all trials was that the inferior–superior component *F*_*y*_ (i.e., the weight-bearing contribution) dominated the load pattern, whereas *F*_*x*_ (posterior–anterior) and *F*_*z*_ (medio– lateral) contributed secondary, task-dependent shear components. The most demanding loading profiles were observed during stairs ascent and fast walking, where peak forces approached ∼5 BW. In contrast, walking, lunge, and stairs descent exhibited comparatively lower magnitudes with values up to 4 BW.

### 3.3 Contact and wear predictions

Among the main outcomes, the task-dependent maps of the contact pressure and the wear depth are presented in Figure 5 and Figure 6, respectively. In addition, the predicted maximum wear depth and volumetric wear values are summarized in Table 2.

**Table 2.**
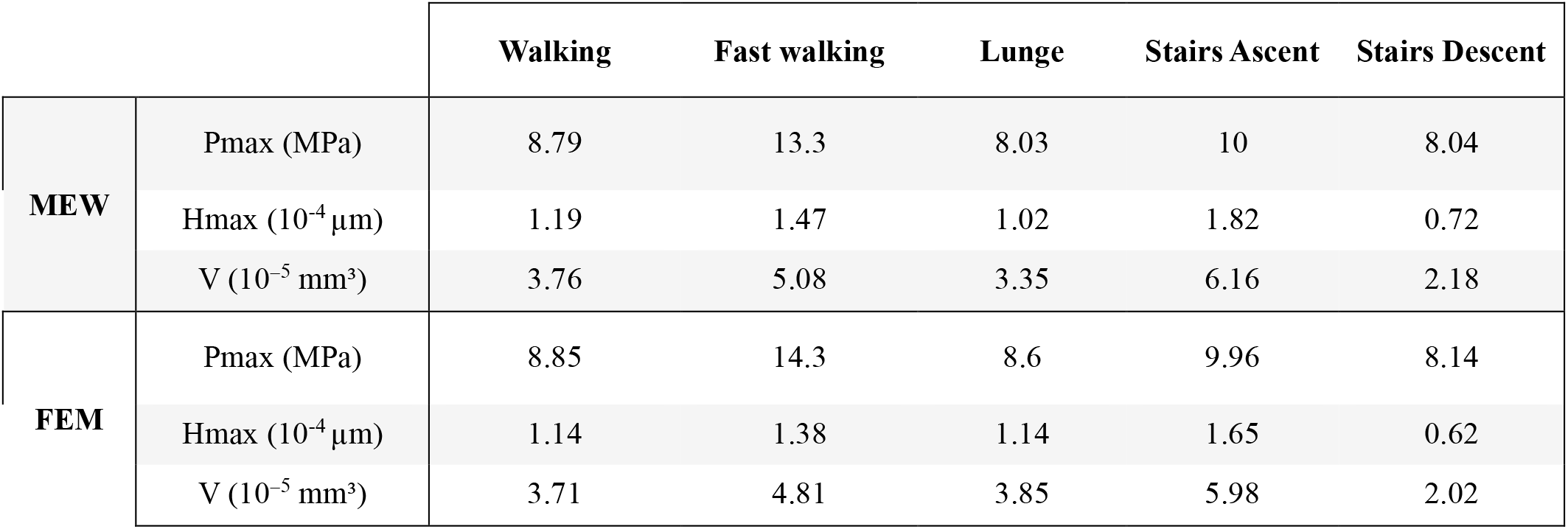
Maximum contact pressure (Pmax), maximum wear depth (Hmax) and wear volume (V) predicted by the MEW method and FEM.

|  |  | Walking | Fast walking | Lunge | Stairs Ascent | Stairs Descent |
| --- | --- | --- | --- | --- | --- | --- |
| MEW | $P_{\max}$ (MPa) | 8.79 | 13.3 | 8.03 | 10 | 8.04 |
| | $H_{\max}$ ($10^{-4} \mu\text{m}$) | 1.19 | 1.47 | 1.02 | 1.82 | 0.72 |
| | $V$ ($10^{-5} \text{ mm}^3$) | 3.76 | 5.08 | 3.35 | 6.16 | 2.18 |
| FEM | $P_{\max}$ (MPa) | 8.85 | 14.3 | 8.6 | 9.96 | 8.14 |
| | $H_{\max}$ ($10^{-4} \mu\text{m}$) | 1.14 | 1.38 | 1.14 | 1.65 | 0.62 |
| | $V$ ($10^{-5} \text{ mm}^3$) | 3.71 | 4.81 | 3.85 | 5.98 | 2.02 |

**Figure 5.**
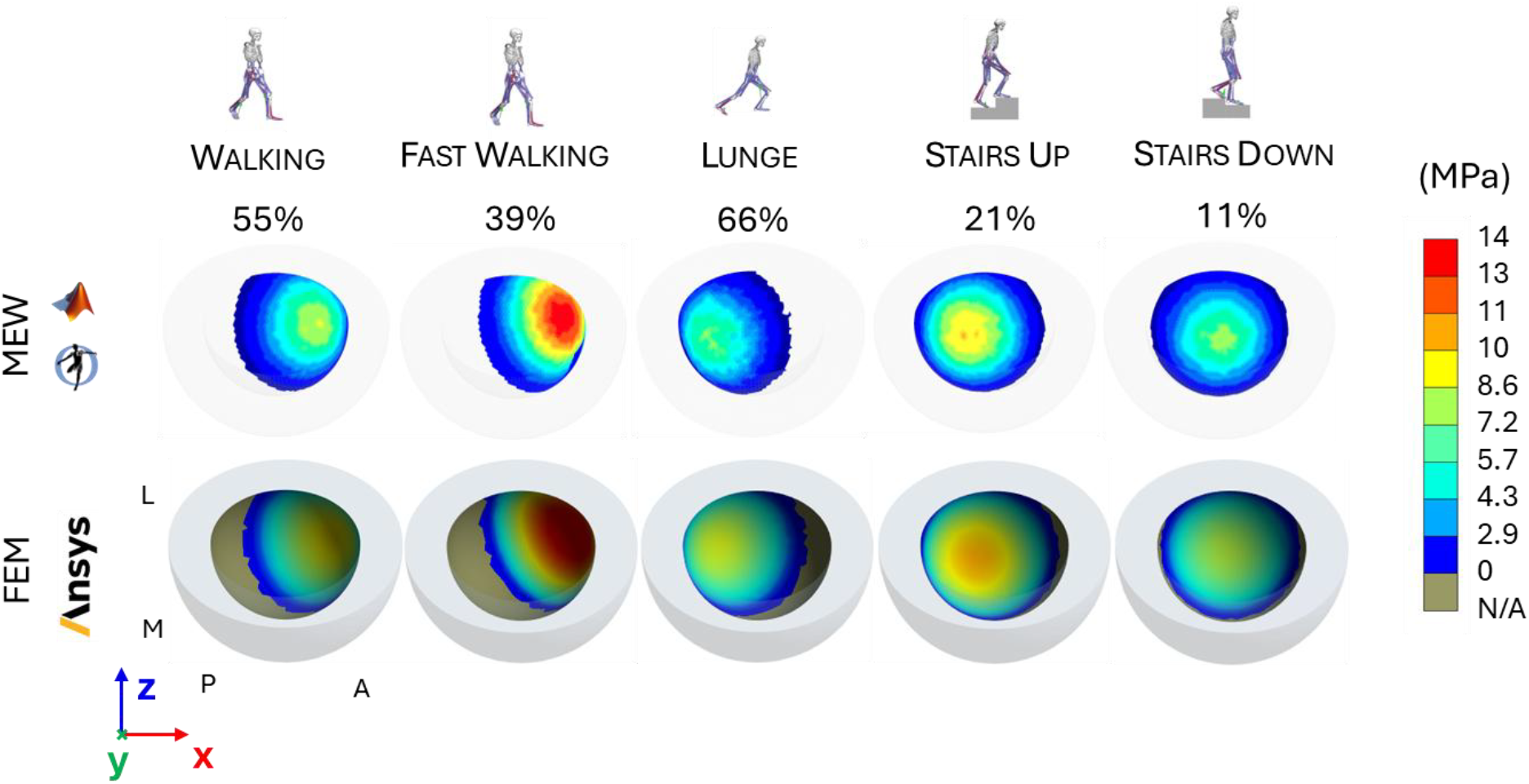
Comparison of pressure maps between the MEW method (top) and FEM (bottom) for the five activities, shown from the same view at each task-specific time point.

**Figure 6.**
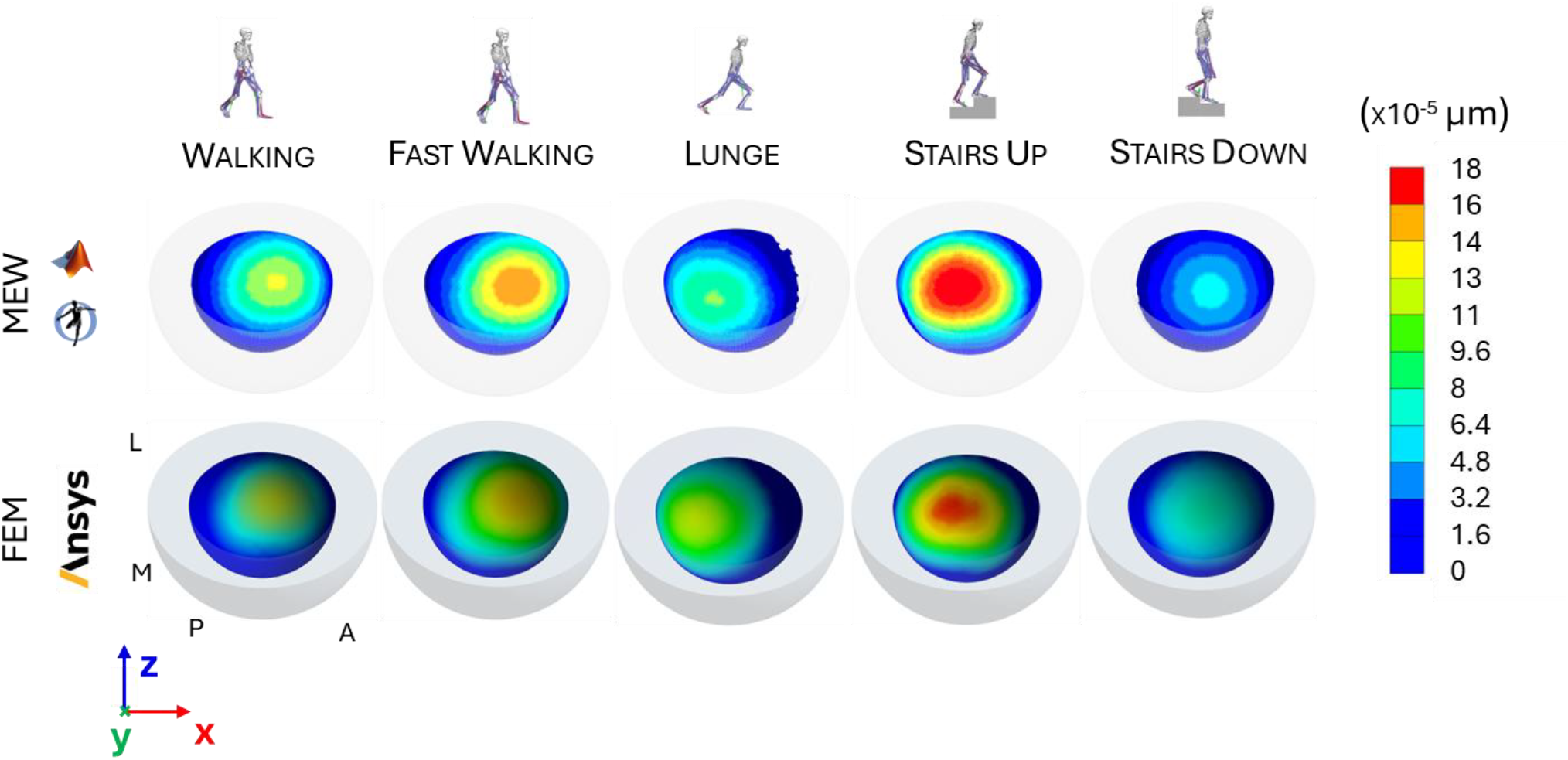
Comparison of wear maps between MEW (top) and FEM (bottom) for the five activities after one task cycle.

**Figure 7.**
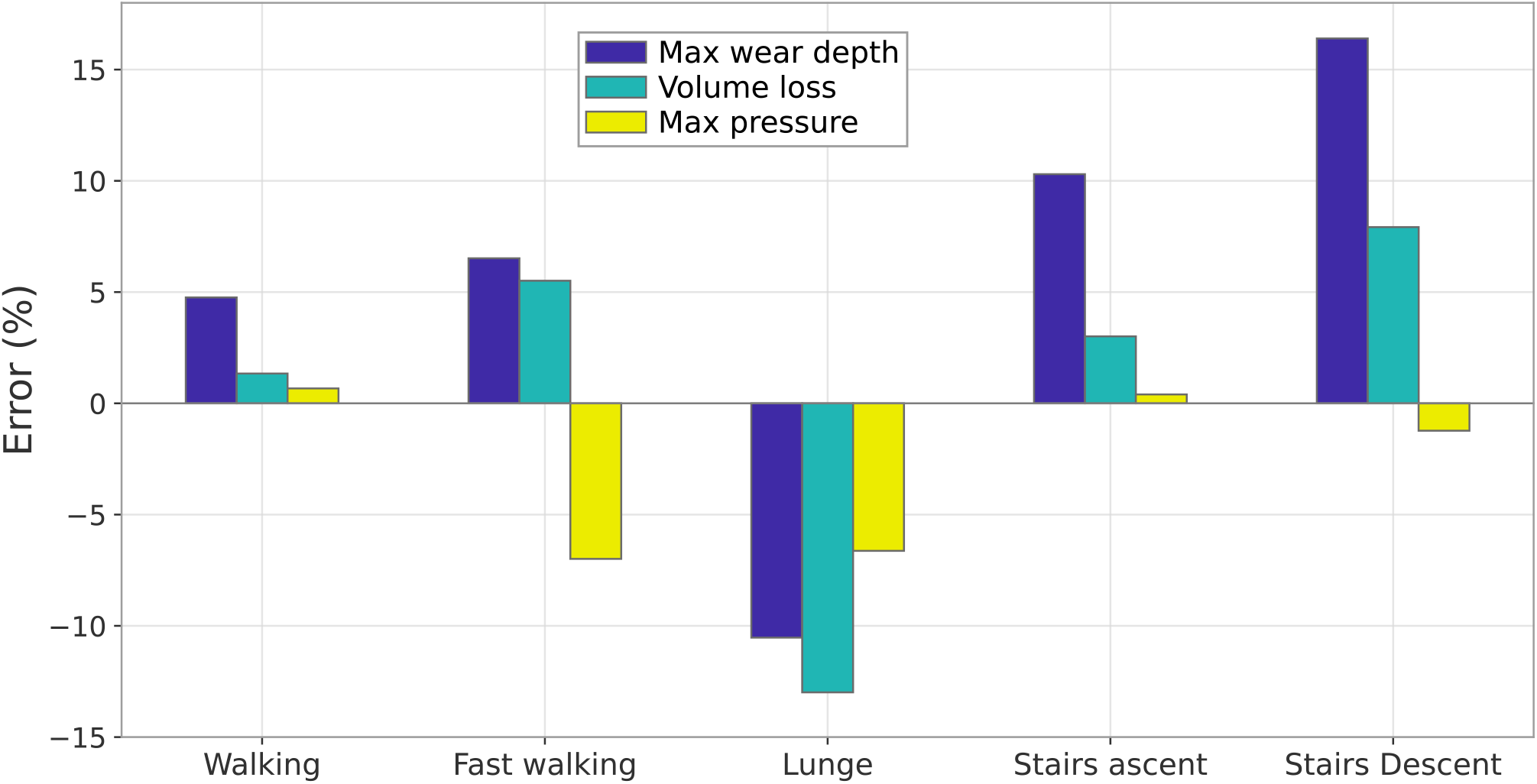
Relative differences in contact and wear parameters between the MEW and FEM methods for each task.

Pressure maps for each motor task (Figure 5) identified the regions of AC subjected to the highest mechanical demand. In most cases, peak contact pressure was in the superior region. However, during fast walking and lunging, the combined effect of joint loading and rigid-body rotation shifted the contact area towards the rim of the AC, anteriorly in fast walking and posteriorly in the lunge, thereby generating the so-called “edge loading” condition.

Among all the tasks, the highest peak contact pressure was predicted during fast walking (13.3 MPa), whereas the lowest was observed during lunge (8.03 MPa) and stairs descent (8.04 MPa). The timing of peak pressure was task-dependent and consistent with the peak longitudinal load component, occurring at 55% for walking, 39% for fast walking, 66% for lunge, 21% for stairs ascent, and 11% for stairs descent.

The wear maps presented in Figure 6 revealed a common wear response across all tasks, whereby the most damaged regions remained confined to the superior area of the AC, consistent with the loading scenario described in Section 3.2.

Linear wear was the highest during stairs ascent (1.82·10^-4^ µm) and fast walking (1.47·10^-4^ µm), followed by walking (1.19·10^-4^ µm) and lunge (1.02·10^-4^ µm), while the lowest value occurred in stairs descent (7.2·10^-5^ µm).

A consistent relationship was also observed between maximum wear depth and total volumetric loss. Tasks associated with greater linear wear, namely stairs ascent and fast walking, produced the largest volumetric losses (6.16·10^−5^ mm^3^ and 5.08·10^−5^ mm^3^, respectively), whereas stairs descent yielded the smallest volume loss (2.18·10^−5^ mm^3^). Walking (3.76·10^−5^ mm^3^) and lunge (3.35·10^−5^ mm^3^) showed intermediate values (Figure 6).

### 3.4 Numerical benchmarking against the finite element method

The spatial patterns and task-dependent trends in the contact and wear maps (Section 3.3; Figures 5 and 6) showed generally limited discrepancies between the MEW and FEM predictions.

With reference to the maximum pressure values reported in Table 2, the differences between the two methods remained within 7%. The largest underestimations were observed during fast walking (– 6.99%) and the lunge (–6.63%), whereas the best agreement was achieved in stairs ascent (+0.4%).

Regarding maximum wear depth, the MEW-obtained results slightly overestimated the FEM-based results for fast walking (+6.52%), stairs ascent (+10.3%), stairs descent (+16.4%), and walking (+4.76%), whereas it underestimated wear in the lunge (–10.53%). For volumetric wear, the framework reproduced benchmark values with small-to-moderate deviations with the same trend of the maximum wear depth: walking +1.34%, fast walking +5.51%, stairs ascent +3.01%, stairs descent +7.92%, and the largest underestimation in the lunge (–12.99%).

### 3.5 Long-term wear simulation

The long-term wear simulation enabled the assessment of the evolution of contact and wear parameters induced by the progressive geometry update.

The evolution of the pressure map at the peak load (i.e. 55% of the walking cycle) is shown in Figure 8, from the unworn configuration up to 4 Mc. Compared with the unworn condition, the maximum contact pressure was initially localized closer to the rim of the AC and progressively evolved towards a more conformal contact pattern, characterized by a reduced peak pressure and a larger contact area. Specifically, as shown in Figure 9a, the maximum contact pressure decreased from 8.8 MPa after the first 0.25 Mc to 7.6 MPa at 4 Mc (Figure 9). A pronounced initial drop was observed immediately after the first geometry update, consistent with a running-in phase associated with an increased wear rate. Subsequently, the pressure followed a non-linear decreasing trend, and the pressure map at 4 Mc qualitatively confirmed an expanded contact area, coherent with the reduction in peak pressure.

**Figure 8.**
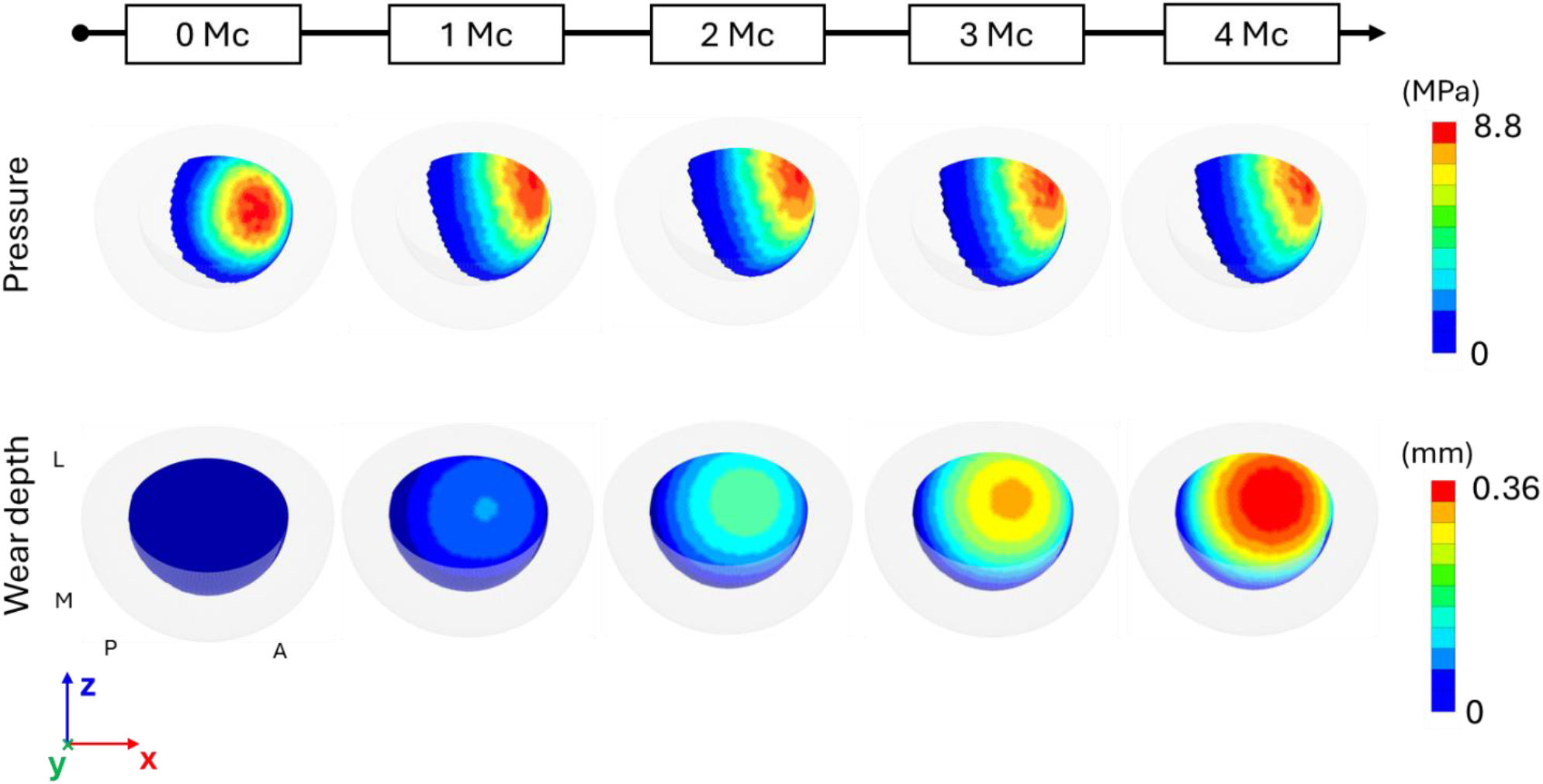
Evolution of contact pressure (top) and wear-depth maps (bottom) over 4 Mc (geometry updated every 0.25 Mc). The contact pressure is represented at the peak load of the walking cycle.

**Figure 9.**
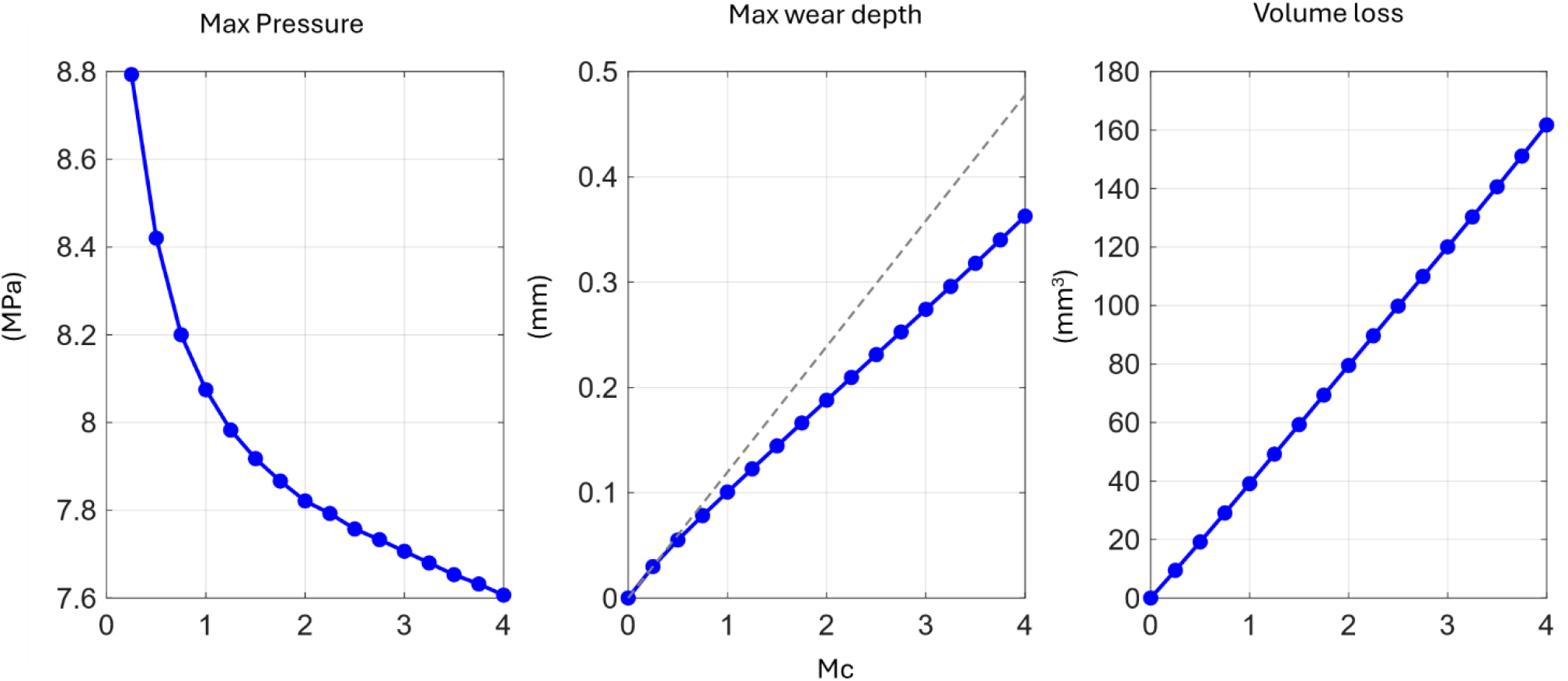
Maximum contact pressure, maximum wear depth and total volume loss over the 4 Mc wear simulation.

The predicted wear, predominantly localized in the anterior region toward the rim of the acetabular cup, was consistent with the forward propulsion phase of gait, as reflected by the direction of the hip joint reaction force during walking (Figure 8).

The maximum linear wear depth increased monotonically but deviated from perfect linearity: an initially steeper slope is followed by a progressive reduction in linear wear rate as geometry updates redistributed contact. At 4 Mc, the maximum wear depth reached 0.363 mm. Compared with the linear extrapolation performed without geometry updating (i.e. case a in sec. 2.3.4, grey dashed line), this corresponded to a reduction of 0.115 mm, indicating that the linear wear approach would overestimate wear approximately of 31%. In contrast, the volumetric wear loss exhibited a trend throughout the simulation, reaching 161 mm^3^ at 4 Mc.

### 3.6 Computational costs

The MEW simulations required, on average, 32 min per second of input kinematics, whereas the corresponding FEM simulations required 1 h 15 min, using an i7 laptop at 2.7 GHz with 16 GB RAM. The long-term walking simulation required 10.5 hours of computational time.

## 4 Discussion and Conclusions

This single-subject proof-of-concept study introduced an MSK-embedded wear (MEW) framework that integrates MATLAB and OpenSim for contact and wear analysis in THA and examined its computational feasibility and numerical consistency with FEM under five daily living conditions.

These last, provided results at two levels: MSK analyses (hip joint kinematics and reaction forces) in OpenSim and contact-wear assessment in MATLAB. With regard to the former, hip angles were obtained with very low kinematic tracking errors during both the CMC and FD simulations, preserving the distinctive features of the subject’s motor-task execution [42]. Similarly, the task-specific hip reaction forces exhibited magnitudes and temporal trends consistent with the literature. In particular, the greater loading demand observed during stair activities and fast walking relative to level walking agrees with previous MSK modelling studies [26,43,44] based on in vivo measurements from instrumented THA implants [9].

Contact and wear assessments were performed using a MATLAB routine and numerically benchmarked against FEM simulations. The differences in contact pressure predicted by the two methods remained within 7%, supporting numerical consistency between the two implementations under the tested conditions. Lunge and fast walking showed the highest edge-loading tendency in this case; this contact condition is known to be more challenging to capture accurately and likely contributed to the increased deviation in pressure prediction. Agreement was also observed for wear predictions, with relative differences in wear depth of up to 16.4% (for stairs descent). Overall, the relative differences in wear volume between the two methods followed those observed for maximum wear depth and remained within ±13%, indicating similar wear-response trends across the analyzed tasks under matched assumptions. The estimated maximum contact pressure during walking was found to be comparable to FEM but also to that reported in other studies [45,46] that considered the EFM under similar loading conditions. These studies evaluated the EFM for other implant geometries and material properties, supporting the plausibility of the present contact predictions but not constituting experimental validation of the MEW workflow.

The EFM mesh-density sensitivity analysis demonstrated that the predicted contact and wear metrics are weakly influenced by surface discretization within the investigated range. Relative to the baseline mesh (6300 elements), variations in maximum wear depth, volumetric loss, and maximum contact pressure consistently remained below 5% for all refinements, indicating effective mesh convergence for the quantities of interest. This limited sensitivity supports the numerical robustness of the proposed method when a uniformly discretized surface mesh is adopted. Conversely, the substantial increase in CPU time with mesh refinement highlights the expected accuracy–computational trade-off and justifies the selection of the baseline mesh as an efficient and reliable discretization for subsequent simulations.

By coupling dynamic and tribological analyses within a single framework, the proposed approach eliminated the need to extract boundary conditions, such as joint reaction forces and rigid-body rotations, from MSK simulations to feed external FEM solvers [17,18,47], ensuring internal consistency between joint loads and wear calculation, and substantially reducing computational effort. This integration provided a unified platform for potentially testing different implant geometries, clearances, and alignment conditions within a single computational workflow, theoretically facilitating design optimization under realistic physiological loading.

Achieving a 4 Mc long-term walking simulation was a relevant outcome, as long-term wear predictions are computationally demanding and often difficult to obtain even within conventional FEM frameworks. In this respect, the proposed workflow enabled prolonged, walking-based simulations at affordable computational cost, making long-term wear assessment practically tractable. The results highlight THA component design and placement as key drivers of functional performance and durability, largely through their influence on edge loading, which can compromise implant stability and ultimately lead to failure [48]. By mapping the expected wear hotspots on the acetabular cup (predominantly anterior in the present case), the method could, after controlled experimental benchmarking and broader multi-subject evaluation, support future investigations of cup orientation and implant sizing. Although direct subject-specific experimental validation was not feasible for the same implant configuration, the predicted volume loss of 160 mm^3^ (40 mm^3^/Mc) was of the same order of magnitude as published values for conventional polyethylene. Parilla et al. reported a median volumetric wear rate of 41.2 mm^3^/year and a median linear wear rate of 0.113 mm/year [49]. However, annual rates are not directly equivalent to per-million-cycle rates without an explicit activity-frequency assumption. Similarly, Ali et al. reported a volumetric wear rate ranging from 30 to 50 mm^3^/Mc for traditional polyethylene acetabular components under in vivo loading conditions [50]. Furthermore, Clarke et al. reported volumetric wear rates for conventional 28 mm UHMWPE acetabular cups of about 32 mm^3^/Mc obtained through experimental hip-simulator tests [51].

Certain limitations should be acknowledged. A main limitation is that, unlike the conventional EFM contact formulation, where penetration is typically evaluated at the centroid of each contact facet, the proposed framework computed it at each vertex of the contact geometry. This choice may introduce numerical discrepancies between the penetrations computed internally by the OpenSim physics engine (Simbody) during the forward dynamics simulation and those kinematically derived by our method; nonetheless, geometry updating was straightforward: it followed directly from the linear Archard wear formulation [10], and directly yielded the updated spatial coordinates defining the worn surface profile.

Second, the analysis was limited to one subject and one representative trial per task, and implant geometry and material properties were assumed because implant-specific information was unavailable. The findings therefore demonstrate workflow feasibility for this case only and cannot establish population-level generalizability, patient-specific predictive accuracy, or clinical utility. Third, both MEW and FEM used a constant wear coefficient and shared key modelling assumptions; consequently, their comparison constitutes numerical benchmarking rather than independent validation. The constant-coefficient assumption also approximates real in vivo THA wear behaviour [12]. Accordingly, this study should be interpreted as a proof of concept for wear simulation and comparative analysis.

Direct subject-specific experimental validation of cumulative in vivo wear was not feasible in this case. Explanted components suitable for quantitative retrieval analysis are not routinely available and no explant was available for the participant considered here. Even if an explant had been available, a strict one-to-one comparison would have remained problematic because the implant’s lifetime loading history cannot be reconstructed: the five laboratory tasks and the single representative trials examined here cannot reproduce the frequency, sequence, intensity, and variability of the activities actually performed by the individual in daily life.

Future work will address these limitations through controlled hip-simulator benchmarking, sensitivity and uncertainty analyses, extension to larger and more heterogeneous cohorts, and adoption of a more mechanistic wear formulation, including wear coefficients that account for cross-shear effects and extending the provided method to metal-on-metal THA implants, where bilateral wear occurs. In addition, long-term simulations will be expanded to mixed-activity protocols that combine multiple daily motor tasks, providing more realistic cumulative wear predictions [14]. Finally, the proposed method could be translated to other joint replacements, such as total knee arthroplasty and shoulder arthroplasty, to support broader implant-level contact and wear investigations.

In conclusion, this single-subject proof of concept demonstrates the feasibility of integrating contact-mechanics and wear assessment within forward musculoskeletal simulations in OpenSim, with numerical consistency relative to FEM under the tested conditions. Beyond wear prediction, the ability to estimate spatially resolved contact variables within the OpenSim workflow may support task-specific assessment of joint contact mechanics and edge loading, as well as evaluation of the effects of implant geometry, clearance, and component positioning.

## Code Availability Statement

The custom MATLAB routines developed for this proof-of-concept study are not publicly available at this stage because the software is under active development and subject to intellectual-property considerations.

