## Supplementary Table 1 for "Embedding wear assessment in musculoskeletal simulation: A proof-of-concept application to total hip arthroplasty"

**Supplementary material**

**Tracking errors of observed kinematics**

Table 3 CMC and FD tracking errors with respect to initial kinematics

|  | **Track. Err. (°)** | **Walk.** | **Fast Walk.** | **Lunge** | **Stairs Up** | **Stairs Down** |
| --- | --- | --- | --- | --- | --- | --- |
| **Hip Fl/Ex** | CMC | -0.1 (0.1) | -0.1 (0.2) | 0 (0.1) | 0 (0.2) | -0.1 (0.1) |
|  | FD | -0.1 (1.1) | 0 (1.6) | 0 (0.4) | 0 (1.2) | -0.3 (0.5) |
| **Hip Ab/Ad** | CMC | 0 (0) | 0 (0.1) | 0 (0) | 0.1 (0.1) | 0.1 (0.1) |
|  | FD | 0 (0.3) | 0 (0.4) | 0 (0.1) | 0.1 (0.4) | 0 (0.3) |
| **Hip Ir/Er** | CMC | 0 (0.1) | 0.2 (0.2) | 0 (0.2) | 0.1 (0.1) | 0 (0) |
|  | FD | 0 (0.5) | 0.2 (0.5) | 0 (0.4) | 0.2 (0.6) | 1 (0.2) |
